# Acoustic analysis and playback experiments do not support the taxonomic revision of the Central and Western Canary Islands subspecies of the Eurasian Stone-curlew (*Burhinus o. distinctus*)

**DOI:** 10.1101/2020.04.09.034249

**Authors:** Marco Dragonetti, Massimo Caprara, Felipe Rodríguez-Godoy, Rubén Barone, V. Rubén Cerdeña, Dimitri Giunchi

## Abstract

**Capsule:** Acoustic analysis does not support the elevation of *B. o. distinctus* to full species.

**Aims:** To verify whether the vocal repertoires of *B. o. oedicnemus* and *B. o. distinctus* show biologically significant quantitative and qualitative differences.

**Methods:** Integration of acoustic analysis of some of the most frequently uttered call types recorded in Italy and in Canary Islands with playback experiments.

**Results:** The vocal repertoires of the individuals belonging to the two subspecies were rather similar, but the quantitative analysis of acoustic parameters evidenced some differences between the considered populations. In particular, the three most used call types showed higher frequency and higher utterance rhythm for *B. o. distinctus* than for *B. o. eodicnemus*. Playback experiments indicated that individuals from the nominate subspecies responded in the same way to the playback of calls of individuals belonging to both subspecies.

**Conclusion:** Acoustic analysis supports the distinctiveness of Stone-curlew populations from Central and Western Canary Islands, thus confirming the available morphological and genetic data. These results, however, do not suggest the elevation of *B. o. distinctus* to full species.

## Introduction

Oceanic islands harbor a species-rich biodiversity and are excellent systems to study evolutionary processes (Mayr *et al*. 2001, Emerson 2002, Grant & Grant 2011). The Canary Islands are an oceanic archipelago consisting of eight principal volcanic islands of varying age (between 1 and 25 Mya) situated about 100-350 km from African mainland and probably never connected to it (Wink 2018). The richness of endemic taxa makes the archipelago one of the most important centres for biodiversity in the temperate region (Juan *et al*. 2000, Illera *et al*. 2012). In relation to extant breeding birds, six species and more than 30 endemic subspecies are currently described (Illera *et al*. 2016).

The Eurasian Stone-curlew *Burhinus oedicnemus* (Stone-curlew hereafter) is a polytypic species with five subspecies described by means of morphological characters distributed in the Palearctic (Cramp & Simmons 1983): *B. o. oedicnemus* (Western and Southern Europe), *B. o. saharae* (Northern Africa and Eastern Mediterranean), *B. o. harterti* (Central Asia), *B. o. insularum* (Eastern Canary Islands) and *B. o. distinctus* (Central and Western Canary Islands). Recent genetic investigations confirmed the four subspecies described for the Western Palearctic and found a significant level of genetic differentiation between Mediterranean and Canary Island populations (Mori *et al*. 2014, 2017). The status of Canarian subspecies is particularly intriguing as several cases of cryptic differentiation of local birds has been discovered in recent years which induces to consider the number of endemic species in the archipelago as probably underestimated (Illera *et al*. 2016, 2018, Sangster *et al*. 2016). Accurate taxonomic designations are of critical importance for biodiversity conservation (Sangster 2018), especially for taxa of a high conservation concern such as *B. o. insularum* and *B. o. distinctus* (Spanish catalogue of threatened species, Real Decreto 139/2011).

Species delineation can be done in many different ways, but the best strategy seems to be the integrative taxonomy, i.e. the combination and integration of multiple types of evidence (Sangster 2018). Together with genetic and morphological information, acoustic data play a major role in birds taxonomic designation, especially for oscine Passerines (see e.g. Illera *et al*. 2014, 2018, Sangster *et al*. 2016, Sangster 2018 and references therein). Indeed, in these species which learn their song structure during the ontogenesis (Mundinger 1982, Podos & Warren 2007), song is considered one of the most important traits promoting differentiation among populations (Price 2008). The repertoire of non-passerines and non-oscine passerines does not include learned vocalizations. However, the calls of a number of these taxa can be considered functionally analogue of the passerine song (Collias 1987, Nuechterlein & Buitron 1998, Galeotti & Sacchi 2001, Lengagne 2001). The geographic variation of vocalizations of non learning species has been much less investigated than that of learning species (Budka *et al*. 2014). A number of study showed however an interesting acoustic variability in non-passerine birds at different geographical scale (e.g. *Gavia immer*, Mager *et al*. 2007; *Crex crex*, Budka et al. 2014, Budka & Osiejuk 2017; *Strix varia*, Odom & Mennill 2012; *Picus viridis*, Fauré 2013). An integrative taxonomic approach including acoustic information turned out to be useful also for non-learning birds (Miller 1983, Robbins & Stiles 1999), allowing the identification as full species of a number of insular taxa (Rheindt *et al*. 2011).

The Stone-curlew is a highly vocal species with a complex and relatively wide vocal repertoire composed of at least 11 different call types (Dragonetti *et al*. 2013), whose geographical variability has never been investigated. In this paper we aim at characterizing the vocalization of two Stone-curlew subspecies, i.e. the nominate and most widespread subspecies and the Canarian subspecies with the smaller population size, *B. o. distinctus* (Delany et al. 2009). In particular we would like to verify whether the well documented morphological and genetic differences of the two subspecies (Cramp & Simmons 1983, Vaughan & Vaughan Jennings 2005, Mori *et al*. 2017) correspond to significant quantitative/qualitative differences in their vocal repertoire. We integrated acoustic analysis of some of the most frequently uttered call types with a preliminary playback experiment aimed at testing whether birds belonging to the nominate subspecies responded differently to calls belonging to their own or to the Canarian subspecies. Playback tests have been successfully used to study song discrimination in passerines (e.g. Tietze *et al*. 2011, Sangster *et al*. 2016), but they have been also employed to examine how subspecies or closely related species of non-passerines respond to calls recorded in different geographical locations (e.g. Curé *et al*. 2010, Miyazaki & Nakagawa 2015). Our final aim is to provide useful data to better assess the taxonomic status of Canarian subspecies, and in particular whether the *distictus* subspecies could be raised to full species status using an integrative taxonomic approach.

## Materials and Methods

### Acoustic analysis

#### Study areas and recording methods

We recorded Stone-curlews vocalizations in Italy, Gran Canaria and Tenerife.

In Italy, in the period 2004-2013, we collected 170 recordings in the Grosseto area (ca. 42°31’ N - 42°51’ N, 11°29’ E - sea coast) throughout the year around. A few recordings (*n* = 8) were made in the Taro River Regional Park (Parma, Italy; 44°74’ N, 10°17’ E) during the reproductive season. All recordings were made using a Fostex FR2 digital recorder, with an Audiotechnica AT815b shotgun microphone or Telinga 22” parabolic dish equipped with a Sennheiser ME62 microphone.

In Gran Canaria, in the year 2015, we collected 47 recordings in various locations (ca. 28°08’ N - 27°59’ N), during the reproductive season. These recordings were made using a TASCAM DR-100 digital recorder with a Sennheiser ME67 shotgun microphone.

Further 30 recordings (some of them with negative results) were made in Tenerife during the reproductive season of the years 2016-2018 in different locations of the southern half of the island (ca. 28°01’ N - 28°20’ N) using a SM2+ Wildlife Acoustic recorder and two omnidirectional SMX-II microphones.

#### Data analysis

We stored all the sound tracks in wav format sampled at 44.1 or 48.0 kHz, with 16-bits accuracy. We used Raven Pro 1.4 (Bioacoustics Research Program 2011) to generate spectrograms, using a Hann window function and DFT size of 1024 samples with 75% overlap. We characterized the vocal repertoire of the three populations by visually comparing the spectrograms of different calls with those published in literature (Cramp & Simmons 1983, Vaughan & Vaughan Jennings 2005, Bergmann *et al*. 2008, Dragonetti *et al*. 2013). We adopted the nomenclature used by Dragonetti *et al*. (2013). We considered each recording as belonging to a different bird and thus the acoustic data were evaluated as independent; it is unlikely that we recorded two or more times the same bird, because in Italy the recordings were made on a wide area and in several years (2004–2013), while in Canary Islands the recordings were made during the reproductive season in different territories. Quantitative analyses were performed on the three most frequently uttered calls: *gallop, kurlee* and *bitonal whistle* (Dragonetti et al. 2013).

##### GALLOP

This call is composed by phrases composed by two/three or more syllables (notes) uttered in rapid succession; many phrases are rhythmically repeated at a near constant time interval, giving way to a call series (or bout) (Figure 1). We determined the number of phrases for each call series, the number of syllables for each phrase, the duration of each series and phrase and then we derived the rhythm of each *gallop* call (phrases/s and syllables/s). Time measurements were automatically done on the oscillograms by setting an amplitude threshold at 10% of the maximal values in Raven Pro. All the available calls were used for these measurements.

**Figure 1.**
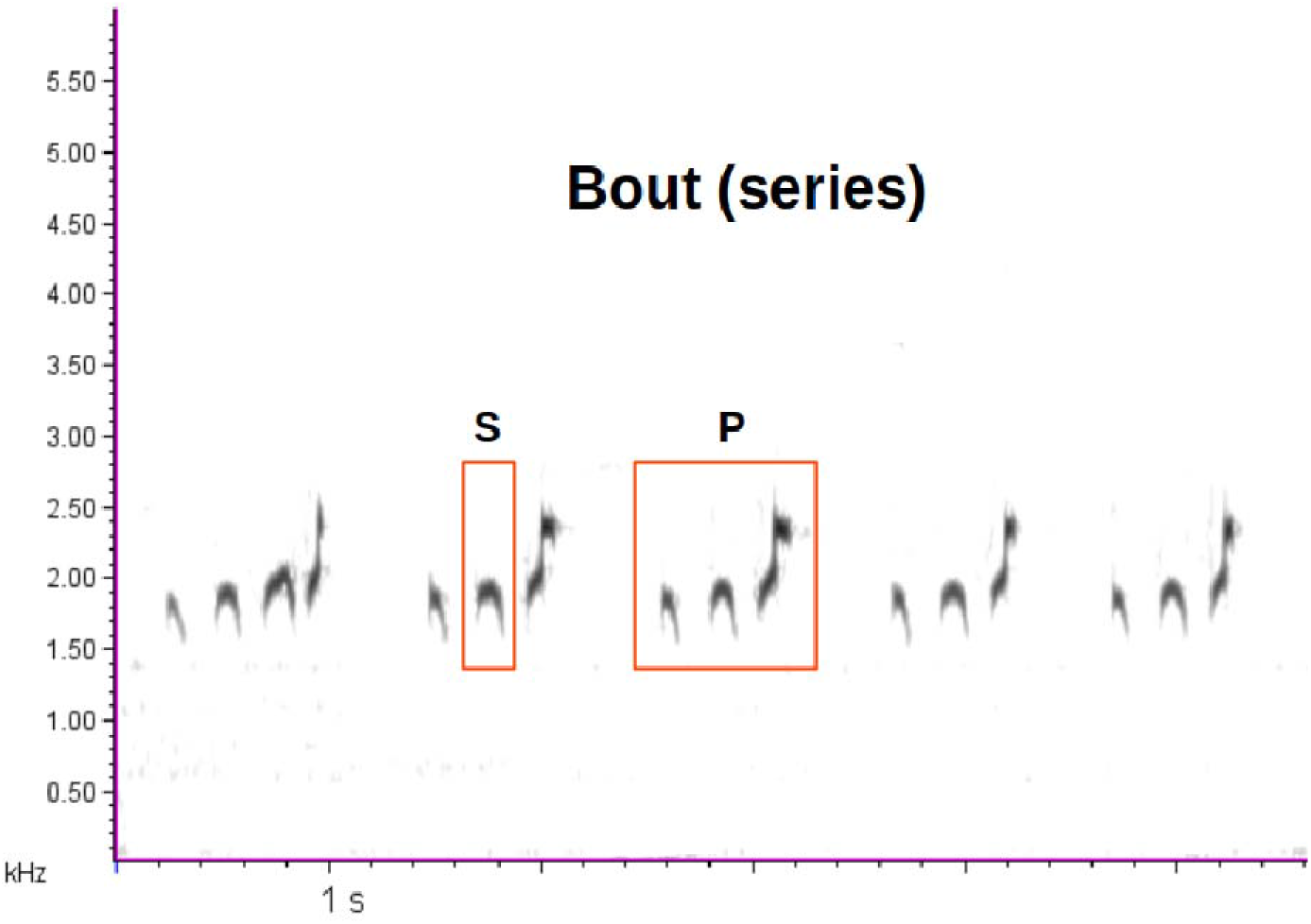
Small part of a *gallop* call bout (series). P = phrase, S = syllable.

Frequency measurements in Hz of the following acoustic parameters were made using the appropriate Raven Pro functions: 1^st^ quartile frequency, 3^rd^ quartile frequency, center frequency. A minimum of four good quality phrases (signal to noise ratio ≥ 2) were randomly selected from each recording and analyzed. When the gallop series was very long (> 16 phrases) we randomly selected 25% of the available phrases.

##### KURLEE

Three good quality calls were randomly selected from each recording and submitted to quantitative analysis. If the available calls were less than three, all calls were analyzed. In the first and the last part of each call (see Figure 2) we measured the 3^rd^ quartile frequency, the peak frequency and the difference between the values recorded in the last and in the first part.

**Figure 2.**
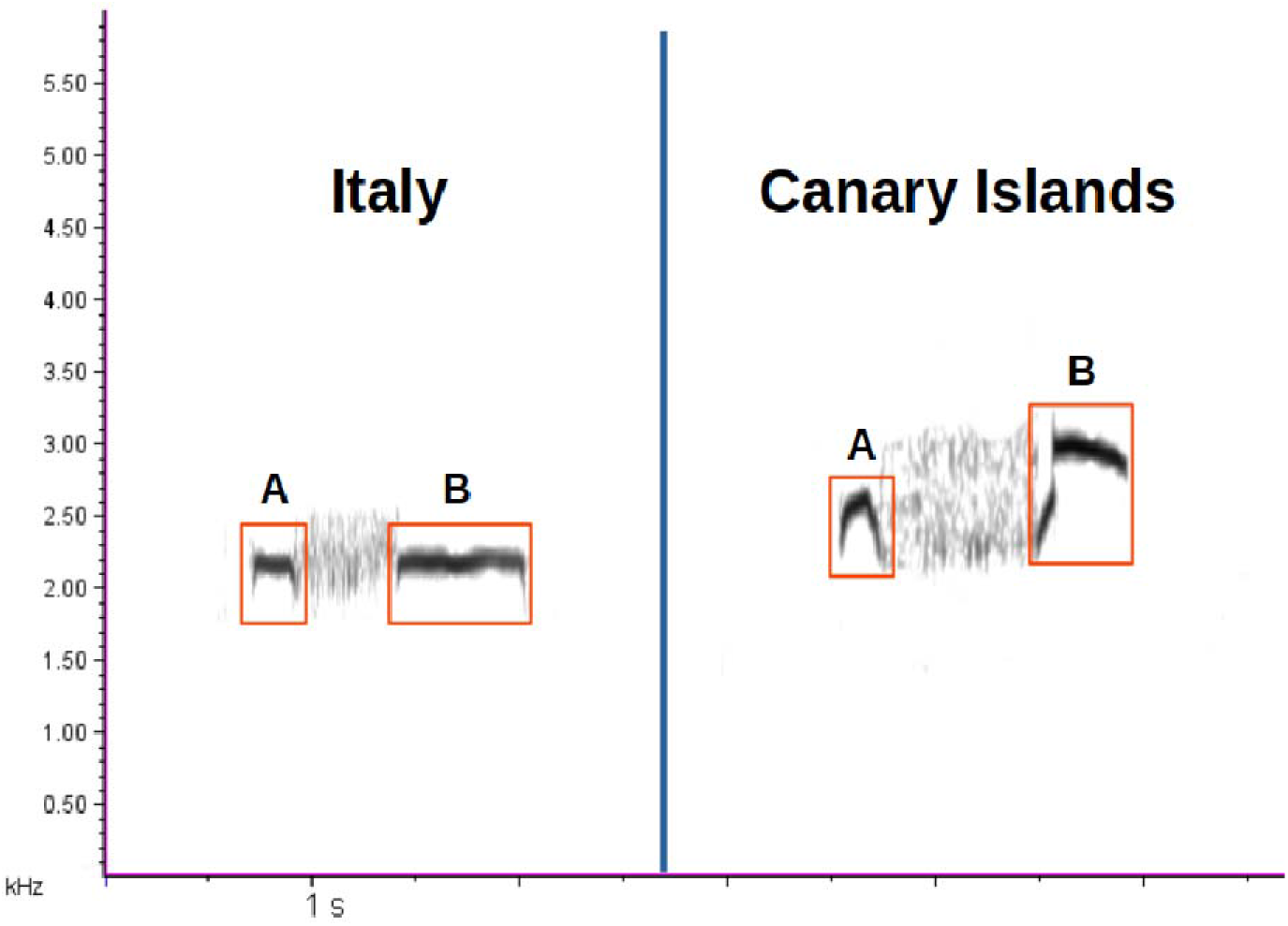
Examples of Canarian and Italian *kurlee* call. Red boxes indicate the call parts that were submitted to quantitative analysis (see Table 2).

##### BITONAL WHISTLE

Four good quality calls (three in some cases, when four good calls were not available) were randomly selected from each recording. We measured peak, center, 1^st^ and 3^rd^ quartile frequency using the appropriate Raven Pro functions. Call duration was measured as described above for *gallop* call.

All the recordings submitted to quantitative analysis were high-pass filtered at 1500 Hz.

#### Statistical analysis

All statistical analyses were performed using R software (R Core Team 2019).

Principal Component Analysis (PCA) was used to reduce the dimensionality of the four considered datasets: *gallop* call (rhythm), *gallop* call (frequency), *kurlee* call, *bitonal whistle*. PCA was calculated on scaled variables using the R-package *FactoMineR* 1.42 (Lê *et al*. 2008). All components with eigenvalue > 1 were considered for each dataset. The second component was extracted even if its eigenvalue was < 1. The retained principal components were compared among populations by means of Linear Mixed Models, considering each recording as random intercept except for the analysis of gallop call rhythm where both recording and bout nested within recording were included as random intercepts. Contrasts were coded to test the following pairwise comparisons: Italy vs. Canary Islands, Gran Canaria vs. Tenerife. The analysis was performed using the R-package *lme4* 1.1.21 (Bates *et al*. 2015). Following Korner-Nievergelt *et al*. (2015), after fitting the model, we simulated 999 values from the joint posterior distribution of the model parameters using the function sim of the R-package *arm* 1.10-1 (Gelman & Hill 2007, Gelman & Su 2018). The 2.5 % and 97.5% quantiles of the simulated values were used as lower and upper limits of the 95% Credible Intervals (95CrI). Model diagnostics and performance (i.e. marginal and conditional R^2^ according to Nakagawa *et al*. (2017) was calculated using the R-package *performance* 0.4.2 (Lüdecke *et al*. 2019).

### Playback experiment

Playback experiments were aimed at testing whether the Italian population of Stone-curlew responded in a different way to the calls recorded in the Canary Islands in comparison to the familiar calls. Tests were carried out in 2018 in the Grosseto area (ca. 42°32’ N −42°52’ N, 11°24’ E - sea coast). Playback tracks were obtained by combining high-quality calls chosen from recordings made in Italy and in Canary Islands as positive stimuli, plus a control stimulus (Nightingale song, recorded in the Grosseto area). We constructed 2-min positive stimuli by combining 30 s *kurlee* call bout + 30 s *gallop* call bout + 20 s *kurlee* call bout + 40 s *gallop* call bout. Since both these call types are usually introduced and/or ended by some *strangled* calls (Dragonetti *et al*. 2013), each *gallop* and *kurlee* bouts included from two to four *strangled* calls, to ensure a more natural sound to playback stimulation. These 2-min samples were digitally filtered out below 1500 Hz. We choose the *gallop* call because it has a reproductive/territorial function and could be probably considered an analogue of the passerine song (Dragonetti *et al*. 2013). The *kurlee* call was chosen because it is the most frequently used call, uttered all year round but mostly during the breeding period (Dragonetti *et al*. 2013). From the 2-min samples (both positive and control stimuli) we prepared the following 56-min playback sequence, which was the same in all experiments:

|  |  |  |  |  |  |  |
| --- | --- | --- | --- | --- | --- | --- |
| Silence | 1<sup>st</sup> stimulus | Silence | 2<sup>nd</sup> stimulus | Silence | 3<sup>rd</sup> stimulus | Silence |
| (20 min) | (2 min) | (10 min) | (2 min) | (10 min) | (2 min) | (10 min) |

The order of presentation of the three stimuli (Stone-curlew from Italy, Stone-curlew from Canary Islands and Nightingale song) was systematically changed in each test. To avoid pseudoreplication we made 12 different tracks using calls of 24 different birds from Canary Islands and 24 from Italy. As Tenerife birds uttered very few and short *gallop* calls (see results below), we could not get good *gallop* samples from this population, therefore only *gallop* calls belonging to birds from Gran Canaria were used.

Playback tracks were uncompressed wav files broadcast using a Samsung smartphone via a Sonic Impact T-Amp external amplifier and a Bird Sound loudspeaker (20-20000 Hz). The amplifier was set to broadcast at an amplitude of 90 dB at 1 m to ensure that playbacks were audible. The amplitude was measured with an AVM2050 (fast response, A-weighting) SPL meter.

We performed the experiments in 29 different locations, chosen on the basis of a preliminary survey aimed at identifying a suitable number of breeding territories in the Grosseto Province. All the experiments begun at sunset (± 5 min) and their timing was controlled by the playback track. The broadcast chain, hidden under a camouflage cloth, was placed near a decoy bird within a breeding territory. One CRENOVA infrared camera (RD1000 Trail Camera) was placed at about 8 m from the decoy to record a video file of the experiment. The camera field of view was about 3 m radius around the decoy and allowed us to detect birds that approached the decoy. At the beginning of the test, one researcher turned on at the same time the broadcasting devices and the infrared camera, and then immediately sat quietly in a hidden place about 30 m from the decoy, pointing a Sennheiser ME67 shotgun microphone connected to a Fostex LE digital recorder towards the decoy in order to record all the vocalizations until the end of the experiment. All recorded tracks were stored in wav format sampled at 44.1 or 48.0 kHz, with 16-bits accuracy.

#### Data analysis

We considered as response only the two call types broadcasted in all playback tracks, i.e. all *gallop* and *kurlee* calls uttered within the first 8 min of silence after each 2 min of playback. Preliminary analyses (not shown) indicated that the vocal response to playback stimulus peaked within 3 min after the end of playback, therefore we have considered only these data in the result section. We considered as an approach response when at least one bird moved at least once toward the decoy to a distance of < 3 m within the first 8 min after each playback.

For each experiment we derived the following parameters: 1) whether at least one bird approached the decoy; 2) total number of *gallop* and *kurlee* calls; 3) total duration of each call type; 4) latency of each call type, that is the time between the beginning of the playback of each 2-min stimulus and the beginning of the first response of each call type. Both *gallop* and *kurlee* calls are often uttered in series. When a given call was uttered alone or alternated with other call types or uttered repeatedly at irregular and long intervals, the duration was the actual call duration and the call number was the actual number of calls; when the same call type was repeated several times at a regular rate leading to a call series, the call duration referred to the entire series (call duration plus silence between calls, which is generally short) and the call number was equal to one.

#### Statistical analysis

Parameters derived from each call type (i.e. latency, total number of calls, total duration of calls) uttered within 3 min after the beginning of the playback of each stimulus were compared among treatments by means of PERMANOVA performed on Euclidean distances calculated on scaled variables (Borcard *et al*. 2018) with 999 permutations, using the function *adonis2* in the R-package *vegan2.5-4* (Oksanen *et al*. 2019). Permutation were constrained within sites to take into account the repeated measures performed on the same site with different treatments. If the result of PERMANOVA was significant, pairwise comparisons were performed using the function *pairwise.adonis2* of the package *pairwiseAdonis* 0.0.1(Martinez Arbizu 2017), adjusting probability according to the False Discovery Rate method (Benjamini & Hochberg 1995).

The effect of the treatment on the likelihood that at least one bird approached the decoy was analysed using a Generalized Linear Mixed Model with a binomial error distribution, with type of playback stimulus as fixed factor and site as random intercept. Contrasts were coded to test the following pairwise comparisons: Italy vs. Canary Islands stimuli, treatments (Italy and Canary Islands) vs control stimuli. Due to failures of the infrared camera, four experiments were not considered in this analysis. The analysis was performed using the R-package *lme4* 1.1.21 (Bates *et al*. 2015) and following the same protocol as reported above for acoustic analysis.

## Results

### Acoustic analysis

The vocal repertoire showed a high similarity between the three populations. In the Canary recordings we have easily recognized 13 out of 15 call types and subtypes described in the Italian repertoire. We did not find only two call types (*answer-to-chick* and *polytonal whistle*), which are uttered in very special circumstances and therefore are less easy to find in ordinary recording sessions (Dragonetti *et al*. 2013). We can reasonably suppose that the vocal repertoire is much the same for Canary and Italian Stone-curlews.

*Kurlee* call and *bitonal whistle* showed minor qualitative differences. Usually *bitonal whistle* is a bitonal syllable, but there is also a less common three tonal version (see Figure 4A in Dragonetti *et al*. 2013), which is very rare for Italian population (3.2% of recorded calls) and more common for Canarian birds (19.7% and 13.0% for Gran Canaria and Tenerife respectively). *Kurlee* call is normally made up by three parts: two whistles interrupted by a rolled or strangled central part. We found that this sequence is doubled very rarely in Italian recordings (1.4%), and much more frequently in Canarian recordings (33.3% and 35.1% for Gran Canaria and Tenerife respectively). Therefore these slight variations of call morphology are not exclusive for the Canarian birds, but are found also in the Italian population, although far less frequently.

Quantitative acoustic analysis showed that the rhythm of *gallop* call differed significantly among populations, as indicated by the PCA analysis (Table 1, Figure S1; PC1: χ^2^ = 5.84, df = 2, p = 0.05, R^2^_marginal_ = 0.93, R^2^_conditionl_ = 0.13; PC2: χ^2^ = 7.16, df = 2, p = 0.03, R^2^_marginal_ = 0.68, R^2^_conditionl_ = 0.10). PC1, which was mainly determined by the number of phrases per bout uttered per second and by phrase duration, tended to be higher in Canary Islands than in Italy (β_Canary Is. vs Italy_ = 0.25, 95CrI = −0.06 – 0.54) and in Gran Canaria than in Tenerife (β_Gran Canaria vs Tenerife_ = 0.49, 95CrI = −0.08 – 1.06). PC2, which was mainly determined by the number of syllables per phrase uttered per second, was not different between Canary Islands and Italy (β_Canary Is. vs Italy_ = 0.07, 95CrI = −0.09 – 0.22), while it was noticeably higher in Gran Canaria than in Tenerife (β_Gran Canaria vs Tenerife_ = 0.34, 95CrI = 0.06 – 0.63). Interestingly, Tenerife birds used *gallop* call far less frequently than the other two populations, as only 133 call phrases were recorded against 809 and 611 for Gran Canaria and Italian birds respectively; also the average length of *gallop* bout was shorter for Tenerife (mean ± SD: 3.9 ± 3.0 s vs 11.0 ± 5.3 s and 11.4 ± 7.5 s for Italy and Gran Canaria respectively).

**Table 1.** *Gallop* call: descriptive statistics (mean ± SD) of the calls recorded in the three considered populations and factor loadings of the acoustic variables on the first two principal components. Eigenvalues and percentage of variance explained by the respective components are given at the bottom of the table. Phrases/s: no. of phrases per bout uttered per second; syllables/s: no. of syllables per phrase uttered per second; phrase duration: phrase length in seconds. In parenthesis, rhythm: no. of recordings, no. of call series, no. of phrases; frequency: no. of recordings, no. of phrases.

| Variable | Population |  |  | PCA |  |
| --- | --- | --- | --- | --- | --- |
|  | Italy | Gran Canaria | Tenerife | PC1 | PC2 |
| Rhythm | (14, 30, 611) | (15, 31, 809) | (9, 16, 133) |  |  |
| Phrases/s | 1.91 ± 0.39 | 2.48 ± 0.67 | 2.35 ± 0.65 | 0.910 | -0.264 |
| Syllables/s | 10.96 ± 1.51 | 13.22 ± 1.91 | 12.61 ± 4.42 | 0.614 | 0.790 |
| Phrase duration (s) | 0.32 ± 0.10 | 0.24 ± 0.10 | 0.23 ± 0.15 | -0.909 | 0.269 |
| Eigenvalue |  |  |  | 2.031 | 0.765 |
| Variance explained |  |  |  | 67.7 | 25.5 |
| Frequency | (15, 82) | (16, 76) | (9, 55) |  |  |
| 1<sup>st</sup> quartile (Hz) | 2318.0 ± 237.8 | 2332.1 ± 220.3 | 2375.0 ± 198.5 | 0.856 | 0.509 |
| Center (Hz) | 2597.0 ± 280.7 | 2715.4 ± 246.4 | 2561.2 ± 214.5 | 0.907 | -0.364 |
| 3<sup>rd</sup> quartile (Hz) | 2470.7 ± 278.8 | 2581.6 ± 269.9 | 2488.1 ± 210.6 | 0.952 | -0.111 |
| Eigenvalue |  |  |  | 2.462 | 0.404 |
| Variance explained |  |  |  | 82.1 | 13.5 |

Frequency parameters of *gallop* call were generally comparable among the three populations (Table 1, Figure S2). Indeed, PC1, which was positively correlated with all variables and explained a large portion of the variance in the sample, did not vary significantly among populations χ^2^ = 1.14, df = 2, p = 0.57, R^2^_marginal_ = 0.67, R^2^_conditionl_ = 0.02). PC2, which contrasted the 1^st^ quartile and the center frequency and explained a marginal part of the variance, was significantly different among populations (χ^2^ = 10.06, df = 2, p = 0.007, R^2^_marginal_ = 0.38, R^2^_conditionl_ = 0.11) and in particular it was comparable between Canary Islands and Italy but noticeably higher in Tenerife than in Gran Canaria (β_Canary Is. vs Italy_ = 0.001, 95CrI = −0.08 – 0.10; β_Gran Canaria vs Tenerife_ = −0.27, 95CrI = −0.4 – −0.09).

*Kurlee* call showed a final part (see Figure 2) with a consistent higher frequency for Canarian than for Italian birds (Table 2, Figure S3). Both principal components differed significantly among populations (PC1: χ^2^ = 36.58, df = 2, p << 0.001, R^2^_marginal_ = 0.62, R^2^_conditionl_ = 0.25; PC2: χ^2^ = 12.22, df = 2, p = 0.002, R^2^_marginal_ = 0.48, R^2^_conditionl_ = 0.08). PC1, which was mainly determined by the frequencies of the second part of the call and by the frequency differences between the second and the first part of the call, was higher in Canary Islands than in Italy (β_Canary Is. vs Italy_ = 0.61, 95CrI = 0.43 – 0.80) but comparable between Gran Canaria and Tenerife (β_Gran Canaria vs Tenerife_ = 0.19, 95CrI = −0.10 – 0.53). PC2, which contrasted the frequencies of the first part of the call with the frequency differences between the second and the first part of the *kurlee*, was not different between Canary Islands and Italy (β_Canary Is. vs Italy_ = 0.06, 95CrI = −0.12 – 0.26), while it was higher in Tenerife than in Gran Canaria (β_Gran Canaria vs Tenerife_ = −0.53, 95CrI = −0.84 – −0.23).

**Table 2.** *Kurlee* call: descriptive statistics (mean ± SD) of the calls recorded in the three considered populations and factor loadings of the six acoustic variables on the first two principal components. Eigenvalues and percentage of variance explained by the respective components are given at the bottom of the table. 3^rd^ quartile A or B: 3^rd^ quartile frequency of the first or the second part of the call (see Fig. 2); Δ quartile: 3^rd^ quartile B - 3^rd^ quartile A; Peak A or B: quartile frequency of the first or the second part of the call; Δ peak: Peak B – Peak A. In parenthesis: no. of recordings, no. of calls.

| Variable | Population |  |  | PCA |  |
| --- | --- | --- | --- | --- | --- |
|  | Italy<br>(27, 81) | Gran Canaria<br>(25, 75) | Tenerife<br>(37, 113) | PC1 | PC2 |
| 3<sup>rd</sup> quartile A (Hz) | 2348.0 ± 207.6 | 2345.7 ± 201.0 | 2482.5 ± 238.4 | 0.227 | 0.930 |
| 3<sup>rd</sup> quartile B (Hz) | 2466.5 ± 255.7 | 2725.7 ± 281.8 | 2686.1 ± 213.1 | 0.902 | 0.224 |
| $\Delta$ quartile (Hz) | 118.5 ± 191.5 | 380.0 ± 308.9 | 203.6 ± 214.3 | 0.738 | -0.589 |
| Peak A (Hz) | 2297.2 ± 184.1 | 2335.4 ± 203.2 | 2455.0 ± 228.1 | 0.239 | 0.927 |
| Peak B (Hz) | 2352.3 ± 254.2 | 2645.4 ± 213.9 | 2665.8 ± 212.7 | 0.900 | 0.249 |
| $\Delta$ peak (Hz) | 55.1 ± 219.7 | 310.0 ± 248.6 | 210.8 ± 215.2 | 0.758 | -0.558 |
| Eigenvalue |  |  |  | 2.850 | 2.494 |
| Variance explained |  |  |  | 47.5 | 41.6 |

For what concerns the variability of frequency characteristics, *bitonal whistle* showed a pattern similar to the *kurlee* call (Table 3, Figure S4). PC1, which was only determined by frequency variables, differed significantly among populations (χ^2^ = 16.84, df = 2, p << 0.001, R^2^_marginal_ = 0.67, R^2^_conditionl_ = 0.21) and it was higher in Canary Islands than in Italy (β_Canary Is. vs Italy_ = 0.56, 95CrI = 0.30 – 0.82) but comparable between Gran Canaria and Tenerife (β_Gran Canaria vs Tenerife_ = 0.34, 95CrI = −0.15 – 0.83). PC2, which mostly represented *bitonal whistle* duration, also differed among populations (χ^2^ = 48.62, df = 2, p << 0.001, R^2^_marginal_ = 0.79, R^2^_conditionl_ = 0.52). In particular, while Canary Islands and Italy resulted to be almost comparable (β_Canary Is. vs Italy_ = −0.06, 95CrI = −0.17 – 0.0), Tenerife showed consistency lower values than Gran Canaria (β_Gran Canaria vs Tenerife_ = 0.86, 95CrI = 0.66 – 1.06).

**Table 3.**
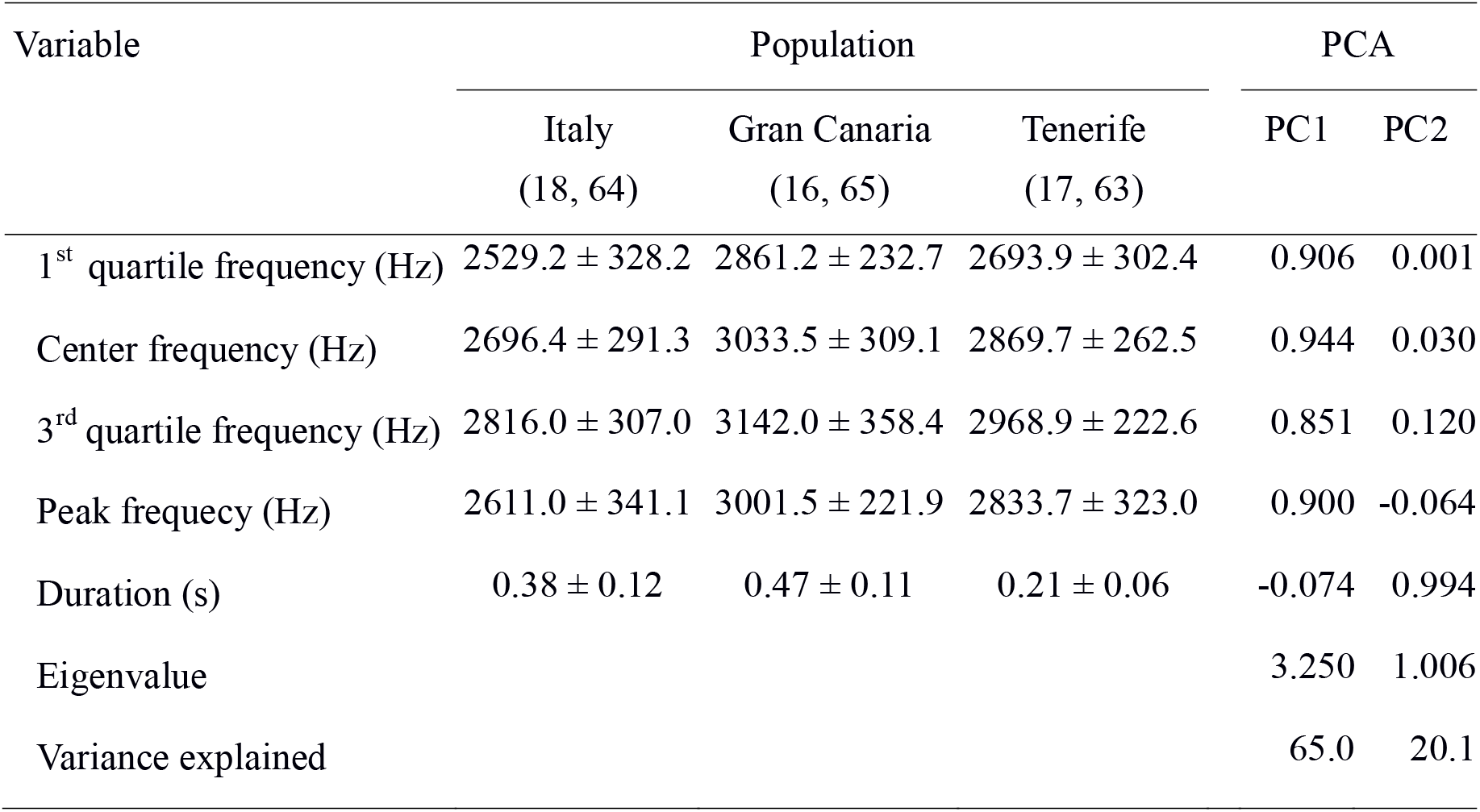
*Bitonal whistle:* descriptive statistics (mean ± SD) of the calls recorded in the three considered populations and factor loadings of the five acoustic variables on the first two principal components. Eigenvalues and percentage of variance explained by the respective components are given at the bottom of the table. In parenthesis: no. of recordings, no. of calls.

| Variable | Population |  |  | PCA |  |
| --- | --- | --- | --- | --- | --- |
|  | Italy<br>(18, 64) | Gran Canaria<br>(16, 65) | Tenerife<br>(17, 63) | PC1 | PC2 |
| 1<sup>st</sup> quartile frequency (Hz) | 2529.2 ± 328.2 | 2861.2 ± 232.7 | 2693.9 ± 302.4 | 0.906 | 0.001 |
| Center frequency (Hz) | 2696.4 ± 291.3 | 3033.5 ± 309.1 | 2869.7 ± 262.5 | 0.944 | 0.030 |
| 3<sup>rd</sup> quartile frequency (Hz) | 2816.0 ± 307.0 | 3142.0 ± 358.4 | 2968.9 ± 222.6 | 0.851 | 0.120 |
| Peak frequency (Hz) | 2611.0 ± 341.1 | 3001.5 ± 221.9 | 2833.7 ± 323.0 | 0.900 | -0.064 |
| Duration (s) | 0.38 ± 0.12 | 0.47 ± 0.11 | 0.21 ± 0.06 | -0.074 | 0.994 |
| Eigenvalue |  |  |  | 3.250 | 1.006 |
| Variance explained |  |  |  | 65.0 | 20.1 |

### Playback experiment

Stone-curlews responded to the playback stimuli with different call types; as stated in the methods section (see above) we studied the responses only considering the two call types (*gallop* and *kurlee* call) used in the playback stimulus.

The descriptive statistics of vocal answers are shown in Table 4. The answers varied significantly among treatments only when considering *gallop* call (*gallop* call: F_2, 84_ = 4.93, p = 0.002; *kurlee* call: F_2, 84_ = 0.81, p = 0.45; PERMANOVA). *Gallop* call emissions did not vary significantly between Italian and Canary Islands stimuli (F_1, 56_ = 0.34, p_adj_ = 0.59; pairwise PERMANOVA), while there was a significant difference between the control and both treatment groups (control vs Italy, F_1, 56_ = 9.51, p_adj_ = 0.002; control vs Canary Islands, F_1, 56_ = 8.21, p_adj_ = 0.002). We observed lower latency, higher number of calls and longer call duration after both treatments with respect to controls (Table 4). It is interesting to observe that the number of syllables/phrase of *gallop* calls uttered after Italian and Canary Islands stimuli was comparable to that of the small number of *gallop* calls uttered during the silence phase at the beginning of the experiments combined with *gallop* calls uttered after the control stimuli (Italian stimuli: 4.2 ± 1.6, *n* = 29; Canary Islands stimuli: 4.3 ± 1.5, *n* = 26; initial silence phase + control stimuli: 4.4 ± 1.2, *n* = 3+4). On the other hand, the *gallop* rhythm (number of syllables/s) tended to be higher in both treatment groups than in controls (Italian stimuli: 12.0 ± 1.7, *n* = 29; Canary Islands stimuli: 12.5 ± 1.8, *n* = 26; initial silence phase + control stimuli: 10.1 ± 2.1, *n* = 3+4).

**Table 4.** Descriptive statistics of the responses to playback stimuli belonging to individuals recorded in Italy or in Canary Islands (control stimulus = nightingale song). Values are mean ± SD in the first 3 min from the beginning of playback of each stimulus (n = 29), except for the last row which reports the number of approaches within 3 m from the decoy (n = 24).

| Response type | Playback stimulus |  |  |
| --- | --- | --- | --- |
|  | Italy | Canary Is. | Control |
| Gallop call |  |  |  |
| Latency (s)<sup>1</sup> | 120.6 ± 70.6 | 138.6 ± 59.2 | 164.0 ± 42.6 |
| Number | 1.0 ± 1.3 | 1.0 ± 1.3 | 0.1 ± 0.4 |
| Duration (s)<sup>2</sup> | 22.8 ± 34.9 | 22.2 ± 35.1 | 1.4 ± 4.2 |
| Kurlee call |  |  |  |
| Latency (s)<sup>1</sup> | 141.5 ± 59.4 | 114.4 ± 69.4 | 123.8 ± 73.5 |
| Number | 1.1 ± 1.7 | 1.6 ± 2.3 | 1.3 ± 1.9 |
| Duration(s)<sup>2</sup> | 8.2 ± 15.6 | 12.9 ± 19.2 | 8.5 ± 12.6 |
| Approaches to the decoy | 8 | 5 | 1 |

<sup>1</sup> Latency was calculated from the beginning of the playback stimulus (latency = 180 s in case of no answer)
<sup>2</sup> Mean duration was calculated by averaging the duration of each call type in all experiments

The probability of approaching the decoy varied significantly among playback stimuli (χ^2^ = 7.84, df = 2, p = 0.02). In particular, we did not record any difference between the two treatments (β_Italy vs Canary Is_. = 0.36, 95CrI = −0.31 – 1.01), while the likelihood of approaching the decoy after the control stimuli was lower than after treatments stimuli (β_Treatments vs Control_ = 0.75, 95CrI = 0.01 – 1.53).

## Discussion

The present work reports the first investigations on population variability of Stone-curlew vocalizations and provides useful information for assessing the taxonomic status of Canarian subspecies.

Our results suggest that the call repertoires of Stone-curlews from Italy and Central-Western Canary Islands are substantially the same, as we could identify almost all call types described in literature in both populations. Two call types, i.e. *kurlee* (the most used call type; Dragonetti *et al*. 2013) and *bitonal whistle* (mainly an alarm signal; Dragonetti *et al*. 2013) show slight morphological differences between Canary Islands and Italy: one subtype of these two calls is by far more common for Canarian birds than for Italian ones, while not being an exclusive character of the insular population. Sharing the same call types and subtypes, but using them in different proportions has been often observed in other sister species (or subspecies) of non-passerine birds, such as Barn Owls (*Tyto a. alba, T. a. schmitzi* and *T. a. detorta*) and Little Owl (*Athene n. noctua* and *A. n. vidalii*) (Robb *et al*. 2015). Fauré (2013) showed that the territorial calls of *Picus viridis* and *P. sharpei* include two syllable types with different dominant frequencies: *P. viridis* uses both types, while *P. sharpei* mostly uses one type only (92% of cases). Interestingly, birds in Tenerife seem to utter gallop call less frequently and with markedly shorter bouts. A possible explanation of this result might be the low density of Stone-curlews in most part of its distribution in this island (Delgado *et al*. 2002, Delany *et al*. 2009, R. Barone & V. R. Cerdeña pers. obs.), which could make the likelihood of active territorial defence and/or disputes relatively low in this island.

The analysis of call morphology and structure emphasized only rather tiny differences, while the quantitative analysis of acoustic parameters provided some clear differences between the considered populations. The gallop call, which is considered a territorial/sexual signal (Dragonetti *et al*. 2013), has an evident higher rhythm in Western Canary Islands and the kurlee call of Canarian birds is characterized by an increase of frequency at its end, while in Italian birds the average frequency remains steady at the end of the call. These are the most evident quantitative differences among populations, but we have also to mention the (slightly) higher frequency parameters of *bitonal whistle* and *gallop* call, which, together with the increase of frequency by the end of *kurlee* call, emphasizethe tendency of Canarian vocalizations to be generally higher pitched. This is probably linked to the smaller body size of this population (Cramp & Simmons 1983, Vaughan & Vaughan Jennings 2005). It is well known, indeed, that the frequencies birds produce are influenced by the anatomy of their vocal tracts (Gaunt & Gaunt 1985, Fletcher & Tarnopolsky 1999), as such frequencies are largely affected by the size and shape of the syrinx, trachea, mouth, and/or bill, which in turn are positively correlated with body size. The negative relationship between frequency, body size, and mass has been observed both within (Podos 2001, Ten Cate *et al*. 2002) and across (Wallschläger 1980, Bertelli & Tubaro 2002) bird species.

The results of playback experiments clearly indicated that Italian birds responded to playback of Canarian and Italian calls in the same way. The multivariate analysis calculated on the latency, number and duration of calls uttered in response to Canarian or Italian playback yielded no significant results and the same outcome was obtained when considering the approach responses to the decoy. The validity of our experimental protocol was confirmed by the significant different responses to both the biological relevant stimuli (Canarian and Italian playback) versus the control stimulus (Nightingale song), which is evident when considering only the *gallop* call and not the *kurlee* call. This result further confirms that *gallop* call is used as a signal of territorial ownership and as an aggressive signal versus a sexual rival (Dragonetti *et al*. 2013). On the other hand, *kurlee* call does not seem to be involved in territorial defence, rather it seems to be a contact call between pair members, neighbours, or adult and young birds. In our experiments, Stone-curlews uttered this vocalization at the same rate after control stimulus and after both treatments, possibly indicating that these calls were simply linked to an increased level of bird activity after sunset, typical of the species (Cramp & Simmons 1983, Caccamo *et al*. 2011).

It is interesting to observe that our data suggest that Stone-curlews increased the rhythm of *gallop* calls signal after a simulated territorial invasion. Indeed, while the average number of syllables/phrase is comparable between *gallop* calls recorded after playback of Stone-curlew calls vs control playback or no playback, the call rhythm is clearly higher after the playback. This result (although preliminary, given the low number of *gallop* calls recorded in control conditions) suggests that in this species the call rhythm is related to the intensity of the intra-specific aggressive behaviour, as observed for other non-passerine birds, such as the Corncrake *Crex crex* (Ręk & Osiejuk 2010), the Junglefowl *Gallus domesticus* (Leonard & Horn 1995) and the American Woodcock *Scolopax minor* (Kroodsma 2005).

To conclude, our results show that the differences between the vocalizations of *B. o. oedicnemus* and *B. o. distinctus* are mostly related to the frequency and rhythm of some calls. Similar variability between different populations was found also in other Charadriiformes, such as *Limnodromus griseus* (Miller *et al*. 1983) and *Charadrius melodus* (Sung *et al*. 2005). The observed differences, however, does not seem big enough for constituting an important prezygotic reproductive barrier. This suggests that local differences in acoustic parameters may not quickly evolve even in isolated populations of non-passerine birds showing almost no migratory behavior, as already observed for some species of songbirds (Illera *et al*. 2014). This is confirmed by the result of the playback experiments, where Italian birds react to acoustic stimuli of both subspecies in the same way.

Our results have to be considered preliminary, because we have no data regarding the vocalizations of birds belonging to the Eastern Canarian and Northwestern African populations. These data would be helpful in order to understand whether the vocal trait variation of Stone-curlews might follow a leapfrog pattern, where populations from opposing ends of a distribution resemble each other in a particular character more than intervening populations (Remsen 1984), as already observed e.g. for doves of the genus *Ptilinopus* (Rheindt *et al*. 2011). Moreover, our playback experiments are incomplete since we could not perform the reciprocal tests in Canary Islands.

Given the above mentioned limitation, the reported acoustic analysis, together with the available morphological (Cramp & Simmons 1983, Vaughan & Vaughan Jennings 2005) and genetic (Mori *et al*. 2014, 2017) data, support the distinctiveness of Stone-curlew populations from Western and Central Canary Islands and further emphasize their conservation importance. The available evidence, however, does not seem enough for suggesting the elevation of *B. o. distinctus* to full species.

One question remain to be answered: what is the cause of the vocal difference between these two subspecies? Regarding the higher pitch of some calls, as stated above, the smaller body size of *B. o. distinctus* could play an important role, but the faster rhythm of the *gallop* call does not seem to be easily explained only considering the above-mentioned morphological differences. Genetic drift on vocal characters should be also taken into account, but it should be considered also that our playback experiment suggest that a faster *gallop* rhythm indicates an increased aggressive attitude and a stronger territorial signal. It might be hypothesized that the faster rhythm of Canary Islands *gallop* calls might be due also to a stronger territorial competition in the area, possibly caused by a relatively restricted area suitable for nesting in the islands with respect to the Italian peninsula. Further research is needed to clarify the taxonomic status of this species and to investigate the significance of the vocal differences in allopatric populations of non-passerine birds.

## Acknowledgments

We are grateful to all the students and volunteers who helped us in the field, and in particular to Valentina Falchi, Luca Passalacqua and Angela Picciau. Thanks are due also to Keith W. Emmerson (†), José J. Hernández and Rayco Jorge, for providing detailed information on some locations appropriated to obtain sound recordings of Stone-curlews in the south of Tenerife Island, and also to Joaquín Vizcaíno (†) for his company during some of the sound recordings.

Examples of the vocalizations described in this paper can be accessed at: http://www.birdsongs.it/songs/burhinus_oedicnemus/burhinus_oedicnemus.html

## References

Bates, D., Mächler, M., Bolker, B. & Walker, S. 2015. Fitting linear mixed-effects models using lme4. J. Stat. Softw. 67: 1–48.

Benjamini, Y. & Hochberg, Y. 1995. Controlling the false discovery rate - A practical and powerful approach to multiple testing. J. R. Stat. Soc. Ser. B-Methodol. 57: 289–300.

Bergmann, H.-H., Helb, H.-W. & Baumann, S. 2008. Die Stimmen Der Vögel Europas. Aula-Verlag, Wiebelsheim, Hunsrück.

Bertelli, S. & Tubaro, P.L. 2002. Body mass and habitat correlates of song structure in a primitive group of birds. Biol. J. Linn. Soc. 77: 423–430.

Bioacoustics Research Program 2011. Raven Pro: Interactive Sound Analysis Software. Version 1.5. The Cornell Lab of Ornithology, Ithaca (NY).

Borcard, D., Gillet, F. & Legendre, P. 2018. Numerical Ecology with R. Springer, Cham.

Budka, M., Mikkelsen, G., Turčoková, L., Fourcade, Y., Dale, S. & Osiejuk, T.S. 2014. Macrogeographic variation in the call of the corncrake *Crex crex*. J. Avian Biol. 45: 65–74.

Budka, M. & Osiejuk, T.S. 2017. Microgeographic call variation in a non-learning species, the Corncrake (*Crex crex*). J. Ornithol. 158: 651–658.

Caccamo, C., Pollonara, E., Emilio Baldaccini, N. & Giunchi, D. 2011. Diurnal and nocturnal ranging behaviour of Stone-curlews *Burhinus oedicnemus* nesting in river habitat. Ibis 153: 707–720.

Collias, N.E. 1987. The vocal repertoire of the red junglefowl: a spectrographic classification and the code of communication. The Condor 89: 510–524.

Cramp, S. & Simmons, K.E.L. 1983. The Birds of the Western Palearctic, Vol. 3. Oxford University Press, London & New York.

Curé, C., Aubin, T. & Mathevon, N. 2010. Intra-sex vocal interactions in two hybridizing seabird species (*Puffinus* sp.). Behav. Ecol. Sociobiol. 64: 1823–1837.

Delany, S., Scott, D., Dodman, T. & Stroud, D. (eds.) 2009. An Atlas of Wader Populations in Africa and Western Eurasia. Wetlands International, Wageningen, The Netherlands.

Delgado, G., Naranjo, J.J., Barone, R., Trujillo, D. & Rodriguez, F. 2002. Datos sobre la distribución de aves esteparias en Tenerife y Gran Canaria, islas Canarias. Vieraea 30: 177–194.

Dragonetti, M., Caccamo, C., Corsi, F., Farsi, F., Giovacchini, P., Pollonara, E. & Giunchi, D. 2013. The Vocal Repertoire of the Eurasian Stone-Curlew (*Burhinus oedicnemus*). Wilson J. Ornithol. 125: 34–49.

Emerson, B.C. 2002. Evolution on oceanic islands: molecular phylogenetic approaches to understanding pattern and process. Mol. Ecol. 11: 951–966.

Fauré, C. 2013. Étude et comparaison des chants du pic vert *Picus viridis viridis* dans le sud-ouest de la France et du pic de sharpe *Picus viridis sharpei* dans le nord de l’Espagne. Alauda 81: 209–225.

Fletcher, N.H. & Tarnopolsky, A. 1999. Acoustics of the avian vocal tract. J. Acoust. Soc. Am. J. Acoust. Soc. Am. 105: 35–49.

Galeotti, P. & Sacchi, R. 2001. Turnover of territorial Scops Owls *Otus scops* as estimated by spectrographic analyses of male hoots. J. Avian Biol. 32: 256–262.

Gaunt, A.S. & Gaunt, S.L.L. 1985. Syringeal structure and avian phonation. Curr. Ornithol. 2: 213–245.

Gelman, A. & Hill, J. 2007. Data Analysis Using Regression and Multilevel/Hierarchical Models. Cambridge University Press New York, NY, USA.

Gelman, A. & Su, Y.-S. 2018. Arm: Data Analysis Using Regression and Multilevel/Hierarchical Models. R Package Version 1.9-3.

Grant, P.R. & Grant, B.R. 2011. How and Why Species Multiply: The Radiation of Darwin’s Finches. Princeton University Press.

Illera, J.C., Palmero, A.M., Laiolo, P., Rodríguez, F., Moreno, Á.C. & Navascués, M. 2014. Genetic, Morphological, and Acoustic Evidence Reveals Lack of Diversification in the Colonization Process in an Island Bird. Evolution 68: 2259–2274.

Illera, J.C., Rando, J.C., Richardson, D.S. & Emerson, B.C. 2012. Age, origins and extinctions of the avifauna of Macaronesia: a synthesis of phylogenetic and fossil information. Quat. Sci. Rev. 50: 14–22.

Illera, J.C., Rando, J.C., Rodriguez-Exposito, E., Hernández, M., Claramunt, S. & Martín, A. 2018. Acoustic, genetic, and morphological analyses of the Canarian common chaffinch complex *Fringilla coelebs* ssp. reveals cryptic diversification. J. Avian Biol. 49: doi:10.1111/jav.01885

Illera, J.C., Spurgin, L.G., Rodriguez-Exposito, E., Nogales, M. & Rando, J.C. 2016. What are We Learning about Speciation and Extinction from the Canary Islands? Ardeola 63: 5–23.

Juan, C., Emerson, B.C., Oromí, P. & Hewitt, G.M. 2000. Colonization and diversification: towards a phylogeographic synthesis for the Canary Islands. Trends Ecol. Evol. 15: 104–109.

Korner-Nievergelt, F., Roth, T., Felten, S. von, Guélat, J., Almasi, B. & Korner-Nievergelt, P. 2015. Bayesian Data Analysis in Ecology Using Linear Models with R, BUGS, and STAN; Including Comparisons to Frequentist Statistics. Elsevier/Academic Press, Amsterdam.

Kroodsma, D.E. 2005. The Singing Life of Birds. Houghton Mifflin, Boston.

Lê, S., Josse, J. & Husson, F. 2008. FactoMineR □: An R Package for Multivariate Analysis. J. Stat. Softw. 25: 1–18.

Lengagne, T. 2001. Temporal stability in the individual features in the calls of eagle owls (*Bubo bubo*). Behaviour 138: 1407–1419.

Leonard, M.L. & Horn, A.G. 1995. Crowing in relation to status in roosters. Anim. Behav. 49: 1283–1290.

Lüdecke, D., Makowski, D. & Waggoner, P. 2019. performance: Assessment of Regression Models Performance. R package version 0.4.2.

Mager, J.N., Walcott, C. & Evers, D. 2007. Macrogeographic variation in the body size and territorial vocalizations of male common loons (*Gavia immer*). Waterbirds 30: 64–72.

Martinez Arbizu, P. 2017. pairwiseAdonis: Pairwise multilevel comparison using adonis. R package version 0.0 1.

Mayr, E., Diamond, J. & Diamond, J.M. 2001. The Birds of Northern Melanesia: Speciation, Ecology & Biogeography. Oxford University Press, Oxford, New York.

Miller, E.H. 1983. The Structure of Aerial Displays in Three Species of Calidridinae (Scolopacidae). The Auk 100: 440–451.

Miller, E.H., Gunn, W.W.H. & Harris, R.E. 1983. Geographic variation in the aerial song of the Short-billed Dowitcher (Aves, Scolopacidae). Can. J. Zool. 61: 2191–2198.

Miyazaki, M. & Nakagawa, S. 2015. Geographical variation in male calls and the effect on female response in little penguins. Acta Ethologica 18: 227–234.

Mori, A., Baldaccini, N.E., Baratti, M., Caccamo, C., Dessì-Fulgheri, F., Grasso, R., Nouira, S., Ouni, R., Pollonara, E., Rodriguez-Godoy, F., Spena, M.T. & Giunchi, D. 2014. A first assessment of genetic variability in the Eurasian Stone-curlew *Burhinus oedicnemus*. Ibis 156:687–692.

Mori, A., Giunchi, D., Rodríguez-Godoy, F., Grasso, R., Baldaccini, N.E. & Baratti, M. 2017. Multilocus approach reveals an incipient differentiation process in the Stone-curlew, *Burhinus oedicnemus* around the Mediterranean basin. Conserv. Genet. 18: 197–209.

Mundinger, P.C. 1982. Microgeographic and Macrogeographic Variation in Acquired Vocalizations of Birds/Eds. Kroodsma DE, Miller EH New-York: Academic Press.

Nakagawa, S., Johnson, P.C.D. & Schielzeth, H. 2017. The coefficient of determination R^2^ and intra-class correlation coefficient from generalized linear mixed-effects models revisited and expanded. J. R. Soc. Interface 14: 20170213.

Nuechterlein, G.L. & Buitron, D. 1998. Interspecific mate choice by late-courting male western grebes. Behav. Ecol. 9: 313–321.

Odom, K.J. & Mennill, D.J. 2012. Inconsistent geographic variation in the calls and duets of Barred Owls (*Strix varia*) across an area of genetic introgression. The Auk 129: 387–398.

Oksanen, J., Blanchet, F.G., Friendly, M., Kindt, R., Legendre, P., McGlinn, D., Minchin, P.R., O’Hara, R.B., Simpson, G.L., Solymos, P., Stevens, M.H.H., Szoecs, E. & Wagner, H. 2019. vegan: Community Ecology Package. R package version 2.5-4.

Podos, J. 2001. Correlated evolution of morphology and vocal signal structure in Darwin’s finches. Nature 409: 185–187.

Podos, J. & Warren, P.S. 2007. The evolution of geographic variation in birdsong. Adv. Study Behav. 37: 403–458.

Price, T. 2008. Speciation in Birds. Roberts and Company. Greenwood Village, CO.

R Core Team 2019. R: A Language and Environment for Statistical Computing. R Foundation for Statistical Computing, Vienna, Austria.

Ręk, P. & Osiejuk, T.S. 2010. Sophistication and simplicity: conventional communication in a rudimentary system. Behav. Ecol. 21: 1203–1210.

Remsen, J.V. 1984. High Incidence of ‘Leapfrog’ Pattern of Geographic Variation in Andean Birds: Implications for the Speciation Process. Science 224: 171–173.

Rheindt, F.E., Eaton, J.A. & Verbelen, F. 2011. Vocal Trait Evolution in a Geographic Leapfrog Pattern: Speciation in the Maroon-Chinned Fruit Dove (*Ptilinopus subgularis*) Complex from Wallacea. Wilson J. Ornithol. 123: 429–440.

Robb, M.S., Berg, A. van den, Constantine, M., Bosman, C.A.W., Petrus, M., Dick Forsman, René Pop & The Sound Approach 2015. Undiscovered Owls: A Sound Approach Guide. The Sound Approach, Poole, Dorset.

Robbins, M.B. & Stiles, F.G. 1999. A new species of pygmy-owl (Strigidae: Glaucidium) from the Pacific slope of the northern Andes. The Auk 116: 305–315.

Sangster, G. 2018. Integrative Taxonomy of Birds: The Nature and Delimitation of Species. In Tietze, D.T. (ed.) Bird Species, 9–37. Springer International Publishing, Cham.

Sangster, G., Rodríguez-Godoy, F., Roselaar, C.S., Robb, M.S. & Luksenburg, J.A. 2016. Integrative taxonomy reveals Europe’s rarest songbird species, the Gran Canaria blue chaffinch *Fringilla polatzeki*. J. Avian Biol. 47: 159–166.

Sung, H.C., Miller, E.H. & Flemming, S.P. 2005. Breeding vocalizations of the piping plover (*Charadrius melodus*): structure, diversity, and repertoire organization. Can. J. Zool. 83: 579–595.

Ten Cate, C., Slabbekoorn, H. & Ballintijn, M.R. 2002. Birdsong and male—male competition: Causes and consequences of vocal variability in the collared dove (*Streptopelia decaocto*). Adv. Study Behav. 31: 31–75.

Tietze, D.T., Martens, J., Sun, Y.-H., Liu Severinghaus, L. & Päckert, M. 2011. Song evolution in the coal tit *Parus ater*. J. Avian Biol. 42: 214–230.

Vaughan, R. & Vaughan Jennings, N. 2005. The Stone Curlew Burhinus oedicnemus. Isabelline Books, Falmouth.

Wallschläger, D. 1980. Correlation of song frequency and body weight in passerine birds. Experientia 36: 412–412.

Wink, M. 2018. Biodiversity on oceanic islands - evolutionary records of past migration events. Heidelb. Jahrb. Online 3: 119–155.

