## Supplementary Figures for "Acoustic analysis and playback experiments do not support the taxonomic revision of the Central and Western Canary Islands subspecies of the Eurasian Stone-curlew (*Burhinus o. distinctus*)"

**Supplementary material**

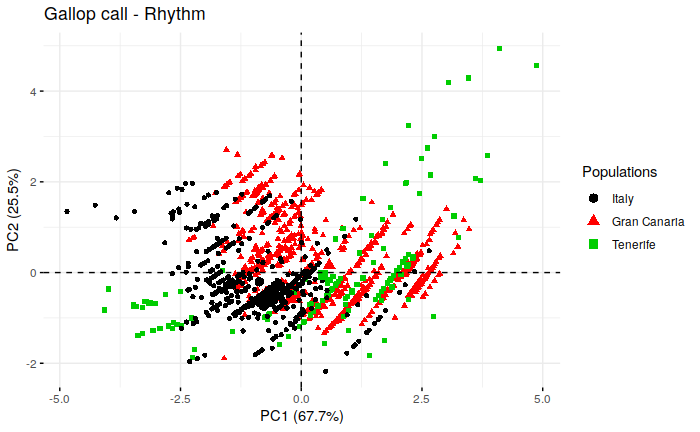

**Figure S1.** Principal component analysis scatterplot of the eigenvectors of the first and second principal components of the three acoustic variables measured for the rhythm of the *gallop* calls recorded in the three considered populations (largest symbols =population centroids). The percentage of variance explained by each component is reported in parentheses.
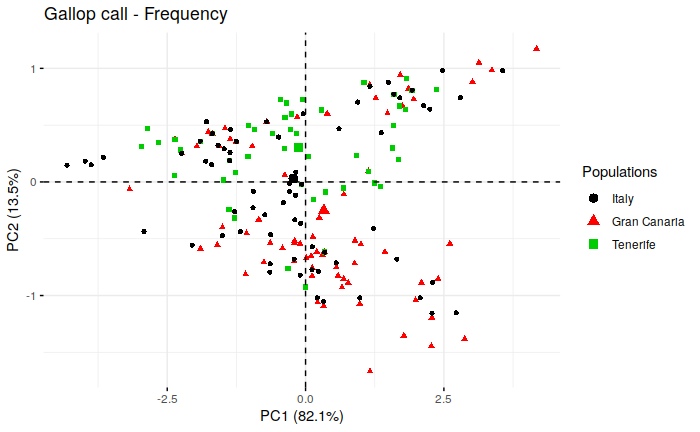

**Figure S2.** Principal component analysis scatterplot of the eigenvectors of the first and second principal components of the three acoustic variables measured for the frequency of the *gallop* calls recorded in the three considered populations (largest symbols =population centroids). The percentage of variance explained by each component is reported in parentheses.

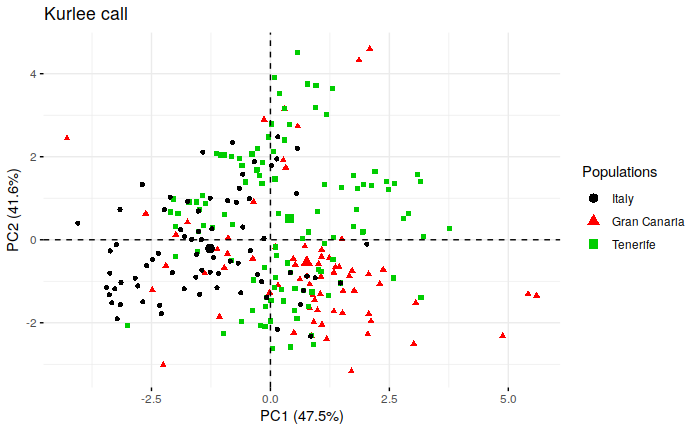

**Figure S3.** Principal component analysis scatterplot of the eigenvectors of the first and second principal components of the six acoustic variables measured for the *kurlee* calls recorded in the three considered populations (largest symbols =population centroids). The percentage of variance explained by each component is reported between parentheses.

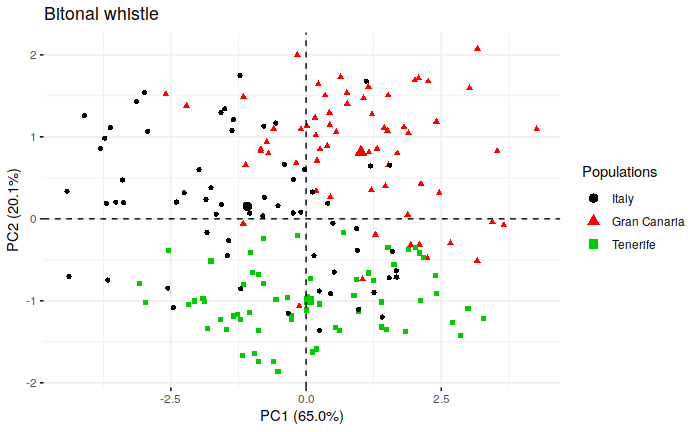

**Figure S4.** Principal component analysis scatterplot of the Eigenvectors of the first and second principal components of the five acoustic variables measured for the *bitonal whistle* calls recorded in the three considered populations (largest symbols =population centroids). The percentage of variance explained by each component is reported between parentheses.
